# Chromosome-level genome assembly of Japanese chestnut (*Castanea crenata* Sieb. et Zucc.) reveals conserved chromosomal segments in woody rosids

**DOI:** 10.1101/2021.07.29.454274

**Authors:** Kenta Shirasawa, Sogo Nishio, Shingo Terakami, Roberto Botta, Daniela Torello Marinoni, Sachiko Isobe

## Abstract

Japanese chestnut (*Castanea crenata* Sieb. et Zucc.), unlike other *Castanea* species, is resistant to most diseases and wasps. However, genomic data of Japanese chestnut that could be used to determine its biotic stress resistance mechanisms have not been reported to date. In this study, we employed long-read sequencing and genetic mapping to generate genome sequences of Japanese chestnut at the chromosome level. Long reads (47.7 Gb; 71.6× genome coverage) were assembled into 781 contigs, with a total length of 721.2 Mb and a contig N50 length of 1.6 Mb. Genome sequences were anchored to the chestnut genetic map, comprising 14,973 single nucleotide polymorphisms (SNPs) and covering 1,807.8 cM map distance, to establish a chromosome-level genome assembly (683.8 Mb), with 69,980 potential protein-encoding genes and 425.5 Mb repetitive sequences. Furthermore, comparative genome structure analysis revealed that Japanese chestnut shares conserved chromosomal segments with woody plants, but not with herbaceous plants, of rosids. Overall, the genome sequence data of Japanese chestnut generated in this study is expected to enhance not only its genetics and genomics but also the evolutionary genomics of woody rosids.

## 1. Introduction

Chestnut is naturally distributed in temperate regions in the northern hemisphere, and plays a critical role in maintaining the landscape and forest ecosystem^1^. Seeds or nuts of chestnut trees provide important nutrition for animals as well as humans, while their wood is used as a source of timber and tannins. The genus *Castanea* (2n = 2× = 24) includes four cultivated species: Japanese chestnut (*C*. *crenata* Sieb. et Zucc), American chestnut (*C*. *dentata* [Marshall] Borkh.), Chinese chestnut (*C*. *mollissima* Bl.), and European chestnut (*C*. *sativa* Mill.). Japanese chestnut is commercially cultivated in Japan and Korea, while the remaining three species are commercially grown in the USA, China, and Europe, respectively. Seguin chestnut (*C*. *seguinii* Dode.) and Henry chestnut (*C*. *henryi* Rehd. et Wils.), found in China, and Allegheny chinkapin (*C*. *pumila* Mill.), which grows in the USA, are recognized as wild species.

In the early 19th century, American chestnut was devastated by chestnut blight, caused by an exotic pathogen, namely, *Cryphonectria parasitica*^2^. European chestnut has also been damaged by chestnut blight and ink disease, caused by *C*. *parasitica* and *Phytophthora cinnamomi*, respectively^3^. In addition, chestnut gall wasp (*Dryocosmus kuriphilus)* was accidentally introduced into Europe, resulting in great damage to chestnut production^4^. Because Japanese chestnut lines as well as Chinese chestnut are generally resistant to diseases and wasps, resistance genes have been transferred from Japanese chestnut to susceptible species in several chestnut breeding programs^3^.

Genome sequence information is useful for not only accelerating breeding programs but also understanding the phylogenetic relationships and revealing the evolutionary history of a given species through comparative genome structure analysis. To the best of our knowledge, genome sequences of three Chinese chestnut lines and one Henry chestnut line have been reported to date^5–8^. However, whole-genome sequence of Japanese chestnut is not yet publicly available. Advances in long-read sequencing technology have enabled the construction of a mega-base-scale genome assembly of haploid, diploid, and polyploid species^9^. Long contig sequences generated by long-read sequencing could be further concatenated at the chromosome level with high-throughput chromosome conformation analyses, optical mapping, and/or genetic mapping to establish pseudomolecule sequences^10^. In this study, using long-read sequencing and genetic mapping, we established chromosome-level pseudomolecule sequences of the Japanese chestnut variety ‘Ginyose’, which has contributed to the spread of chestnut cultivation in Japan owing to its large nut size and excellent appearance. Despite its long history of cultivation that dates back to the 17th century, ‘Ginyose’ is still a major cultivated variety in Japan, accounting for approximately 15% of the national chestnut production. Genetic structure and parentage analysis shows that ‘Ginyose’ is the predecessor of many Japanese cultivars^11^.

## 2. Materials and methods

### 2.1 Plant materials and DNA extraction

A tree of the Japanese chestnut cultivar ‘Ginyose’ planted at the Institute of Fruit Tree and Tea Science, NARO (Tsukuba, Japan) was used for genome sequencing. Genomic DNA was extracted from tree leaves with Genomic-tips (Qiagen, Hilden, Germany).

### 2.2 Genome sequencing and analysis

Software tools used for data analyses are listed in Supplementary Table S1. Genomic DNA libraries for short- and long-read sequencing were generated with the TruSeq DNA PCR-Free Kit (Illumina, San Diego, CA, USA) and the SMRTbell Express Template Prep Kit 2.0 (PacBio, Menlo Park, CA, USA), respectively. Short-read sequence data were obtained using the HiSeq2000 platform (Illumina). Genome size was estimated using Jellyfish after removing adaptor sequences (AGATCGGAAGAGC) and reads obtained from organelle genomes (GenBank accession numbers: HQ336406 and MN199236). Long-read sequence data were obtained using the Sequel system (PacBio), and primary contigs and alternate contigs, which were generated from one allele and the other, respectively, of diploid genomes, were assembled with Falcon. Then, haplotype sequences were resolved to generate haplotigs with Falcon_Unzip. Sequence errors in the contigs were corrected twice using long reads with ARROW, and potential haplotype duplications in the primary contigs were removed with Purge_Dups. Contig sequences potentially contaminated from organelle genomes (GenBank accession numbers: HQ336406 and MN199236), which showed sequence similarity of > 90% with Minimap2, were removed. Assembly completeness was evaluated with the embryophyta_odb10 data using Benchmarking Universal Single-Copy Orthologs (BUSCO).

### 2.3 Genetic map-based chromosome-level sequence construction

Reads obtained by double digest restriction site-associated DNA sequencing (ddRAD-Seq) of an F1 mapping population (n = 185), derived from a cross between ‘Bouche de Bétizac’ (*C*. *sativa* × *C*. *crenata*) and ‘Madonna’ (*C*. *sativa*) ^4^, were mapped onto the primary contigs as a reference using Bowtie2, and sequence variants were detected with BCFtools. High-confidence single nucleotide polymorphisms (SNPs) were selected with VCFtools (parameters of --minDP 5 --minQ 999 --max-missing 0.5 --maf 0.2 --max-maf 0.8) and subjected to linkage analysis with Lep-Map3. The nomenclature and direction of the linkage groups were based on the map reported by Torello Marinoni et al.^4^ To construct pseudomolecule sequences at the chromosome level, primary contigs were assigned to the resultant genetic map with ALLMAPS.

### 2.4 Repeat sequence analysis

Repetitive sequences in the assembly were identified with RepeatMasker using repeat sequences registered in Repbase and a de novo repeat library built with RepeatModeler. Repeat elements were classified into nine types with RepeatMasker: short interspersed nuclear elements (SINEs), long interspersed nuclear elements (LINEs), long terminal repeat (LTR) elements, DNA elements, small RNA, satellites, simple repeats, low complexity repeats, and unclassified.

### 2.5 Gene prediction and annotation

Protein-coding genes were predicted using a MAKER pipeline, based on peptide sequences predicted in the *C*. *mollissima* genome (GWHANWH00000000, *n* = 33,597)^7^ and transcriptome sequences of three *Castanea* species^2^, *C*. *crenata* (cci_ccn, *n* = 13,451), *C*. *dentata* (AC454_v3, *n* = 45,288), and *C*. *sativa* (csi_csn, *n* = 12,771), released in the Hardwood Genomics Project (https://doi.org/10.25504/FAIRsharing.srgkaf). Short gene sequences (< 300 bp), genes in repeat sequences, and genes with annotation edit distance (AED) > 0.5, which is proposed as a threshold for good annotations in the MAKER protocol, were removed to facilitate the selection of high-confidence genes. Functional annotation of the predicted genes was performed with Hayai-Annotation Plants.

### 2.6 Comparative genome structure analysis

The genome structure of Japanese chestnut cultivar ‘Ginyose’ was aligned to the genome sequence of Chinese chestnut (*C*. *mollissima*), as a reference, with Unimap. In addition, publicly available chromosome-level sequences of 114 plants^10^, which, in accordance with Angiosperm Phylogeny Group (APG) classification IV^12^, consisted of 20 monocots, 93 eudicots (including 66 rosids and 23 asterids), and one member of Nymphaea, were also employed as references. Dot-plot charts based on the alignments were drawn with D-Genies.

## 3. Results and data description

### 3.1 Genome sequence and assembly

In accordance with k-mer frequency analysis of 63.3 Gb short reads, the genome size of *C*. *crenata* was estimated at 665.9 Mb (Figure 1). A total of 2.7 million long reads (47.7 Gb, 71.6× genome coverage, N50 = 29.8 kb) obtained from four single-molecule real-time (SMRT) cells were assembled into two haplotype-resolved sequences: primary contigs and haplotigs. The resultant assemblies of primary contigs (CCR_r1.0) and haplotigs (CCR_r1.0.haplotigs) consisted of 781 and 3,355 sequences, respectively (Table 1). Total lengths of primary contigs (CCR_r1.0) and haplotigs (CCR_r1.0.haplotigs) were 721 Mb (N50 = 1.6 Mb) and 629.2 Mb (N50 = 275.8 kb), respectively. The completeness of primary contigs indicated that 96.6% of sequences were complete BUSCOs (Table 1).

**Figure 1.**
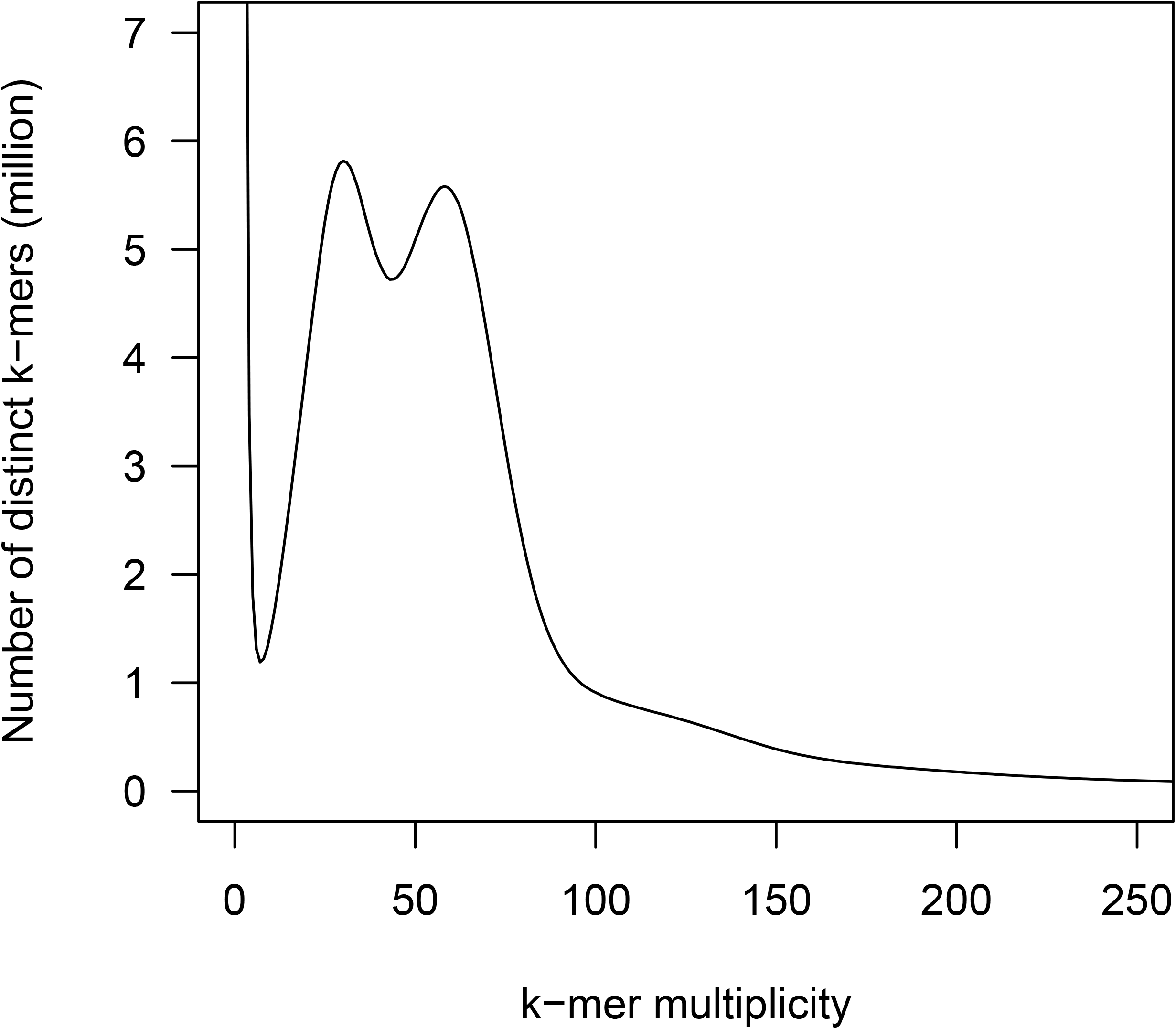
Estimation of the genome size of Japanese chestnut (*Castanea crenata*), based on *k*-mer analysis (*k* = 17) with the given multiplicity values.

**Table 1.** Statistics of the contig sequences of Japanese chestnut

|  | CCR_r1.0 | CCR_r1.0.haplotig |
| --- | --- | --- |
| Total contig size (bases) | 721,168,657 | 629,166,897 |
| Number of contigs | 781 | 3,355 |
| Contig N50 length (bases) | 1,595,543 | 275,768 |
| Longest contig size (bases) | 13,981,633 | 1,688,135 |
| Gap (bases) | 0 | 0 |
| Complete BUSCOs | 96.6 | n.a. |
| Single-copy BUSCOs | 91.9 | n.a. |
| Duplicated BUSCOs | 4.7 | n.a. |
| Fragmented BUSCOs | 0.9 | n.a. |
| Missing BUSCOs | 2.5 | n.a. |
| #Genes | 69,980 | n.a. |

2 n.a.: not analyzed.

### 3.2 Pseudomolecule sequence construction

To construct a genetic map of *C*. *crenata*, we used ddRAD-Seq data of a mapping population reported prreviously^4^. A total of 903 million high-quality ddRAD-Seq reads obtained from the mapping population and parental lines were mapped onto the primary contigs, with an average mapping rate of 86.5%. SNPs were detected and filtered. Consequently, 15,364 high-quality SNPs were identified, of which 14,973 were grouped into 12 linkage groups, which corresponds to the number of chromosomes in *C*. *crenata*. Then, SNP order and the map distance between adjacent SNPs were calculated (Table 2, Supplementary Table S2). The nomenclature of linkage groups was based on our previous study^4^. A total of 575 primary contigs (683.8 Mb) were assigned to this genetic map and connected with 100 Ns to establish pseudomolecule sequences, while the remaining 206 contigs (37.4 Mb) were not assigned to any linkage groups (Table 2).

**Table 2.** Statistics of Japanese chestnut pseudomolecule sequences (CCR_r1.0.pmol)

| Linkage<br>group | No. of<br>SNPs | Genetic<br>distance<br>(cM) | No. of<br>bins | No. of<br>contigs | % | Total length (bp) | % |
| --- | --- | --- | --- | --- | --- | --- | --- |
| CCR1.0A | 2,483 | 173.7 | 284 | 59 | 7.6 | 89,228,932 | 12.4 |
| CCR1.0B | 1,514 | 172.2 | 202 | 53 | 6.8 | 53,118,407 | 7.4 |
| CCR1.0C | 1,232 | 152.6 | 204 | 44 | 5.6 | 56,655,719 | 7.9 |
| CCR1.0D | 1,083 | 137.5 | 161 | 57 | 7.3 | 52,875,950 | 7.3 |
| CCR1.0E | 1,342 | 165.2 | 228 | 61 | 7.8 | 71,160,265 | 9.9 |
| CCR1.0F | 1,339 | 132.3 | 154 | 42 | 5.4 | 61,824,531 | 8.6 |
| CCR1.0G | 924 | 137.2 | 153 | 44 | 5.6 | 44,683,286 | 6.2 |
| CCR1.0H | 816 | 138.9 | 170 | 34 | 4.4 | 53,252,922 | 7.4 |
| CCR1.0I | 1,199 | 157.9 | 168 | 42 | 5.4 | 45,426,588 | 6.3 |
| CCR1.0J | 1,036 | 148 | 157 | 32 | 4.1 | 40,655,448 | 5.6 |
| CCR1.0K | 1,026 | 144.9 | 196 | 62 | 7.9 | 67,434,223 | 9.4 |
| CCR1.0L | 979 | 147.4 | 140 | 45 | 5.8 | 47,439,869 | 6.6 |
| **Total** | **14,973** | **1,808** | **2,217** | **575** | **73.6** | **683,756,140** | **94.8** |

### 3.3 Repeat sequence analysis and gene prediction

Repetitive sequences (425.5 Mb) accounted for 59.0% of the Japanese chestnut genome assembly (Table 3). LTR retroelements were the most abundant repeat sequences (172.7 Mb, 24.0%), followed by unclassified repeat sequences (154.0 Mb, 21.3%), i.e., those unavailable in public databases.

**Table 3.** Repetitive sequences in the Japanese chestnut genome

| Repeat type | No. of elements | Physical length<br>(bp) | % |
| --- | --- | --- | --- |
| SINEs | 3,222 | 349,743 | 0.0 |
| LINEs | 58,824 | 28,244,400 | 3.9 |
| LTR elements | 173,629 | 172,741,932 | 24.0 |
| DNA transposons | 206,227 | 47,812,722 | 6.6 |
| Unclassified | 672,305 | 153,965,018 | 21.3 |
| Small RNA | 1,548 | 851,212 | 0.1 |
| Satellites | 1,888 | 1,522,847 | 0.2 |
| Simple repeats | 308,508 | 12,661,150 | 1.8 |
| Low complexity | 43,949 | 2,095,795 | 0.3 |

A total of 195,950 potential protein-coding sequences were identified in the Japanese chestnut genome assembly, based on *ab initio* prediction and amino acid sequence homology among four *Castanea* species: *C*. *crenata*, *C*. *dentata*, *C*. *mollissima*, and *C*. *sativa*. Subsequently, protein-coding sequences were filtered to remove 89,194 genes with AED > 0.5, 13,389 protein-coding sequences in repeat sequences, 23,386 short sequences (< 300 bp), and one redundant sequence. Finally, 69,980 sequences were identified as high-confidence genes, including 83.6% complete BUSCOs. Functional annotation analysis showed that of the 69,980 predicted genes, 4,370, 7,803, and 5,219 sequences were assigned to Gene Ontology slim terms in the biological process, cellular component, and molecular function categories, respectively, and 1,267 genes had enzyme commission numbers (Supplementary Table S3).

Next, peptide sequences of *C*. *mollissima* and transcriptome sequences of *C*. *crenata*, *C*. *dentata*, and *C*. *sativa* were aligned to the genome sequence of Japanese chestnut obtained in this study. A total of 24,561 genes predicted in the Japanese chestnut genome sequence were covered by transcriptome sequences from at least one of the three *Castanea* species (Figure 2), while the remaining 28,773 genes did not correspond to transcriptome sequences from any *Castanea* species.

**Figure 2.**
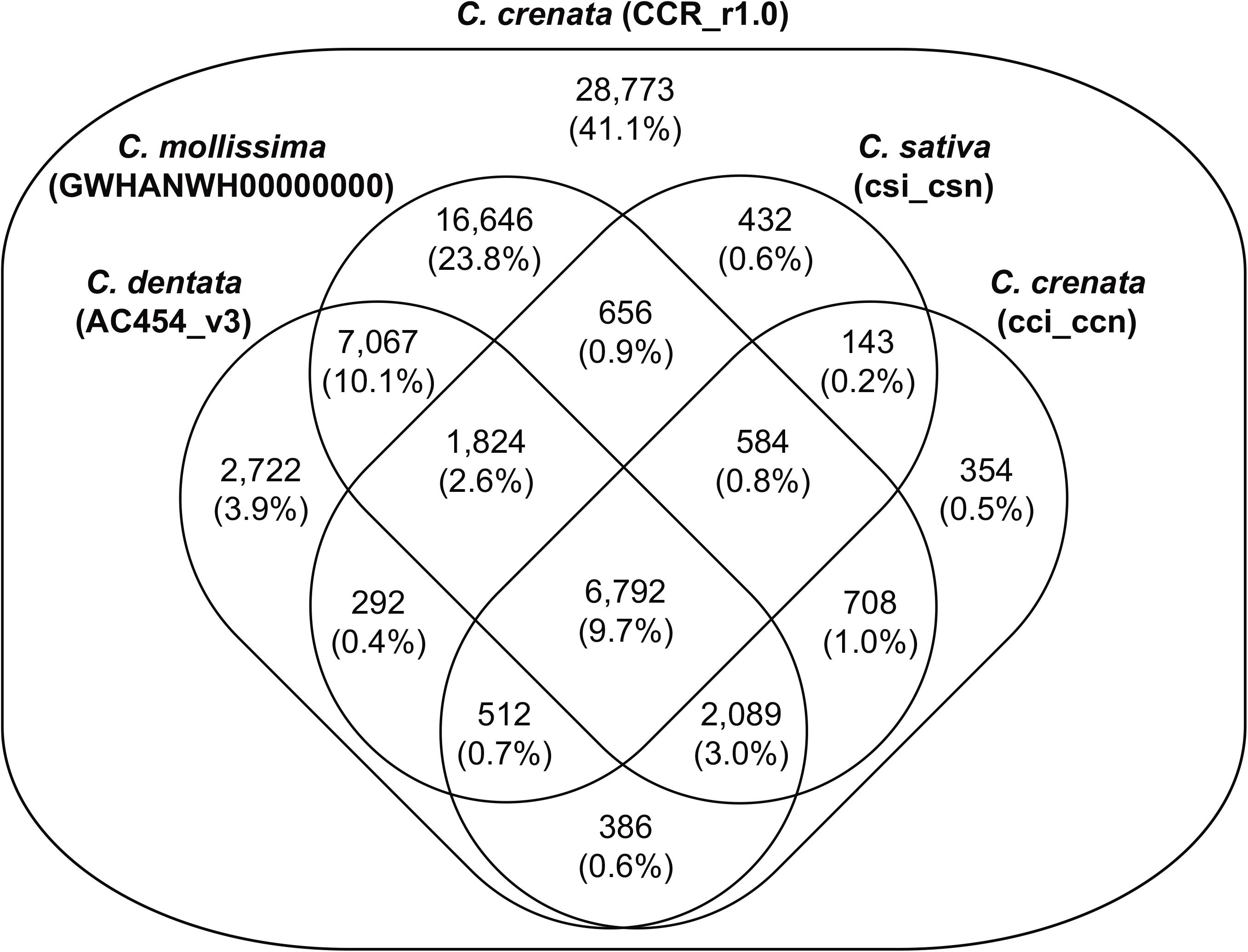
Number of genes in the genomes of Japanese chestnut and Chinese chestnut supported by members of the Rosaceae family. Predicted genes in the genomes of Japanese chestnut (CCR_r1.0) and Chinese chestnut (GWHANWH00000000), and the transcriptome sequences of American chestnut (AC454_v3), European chestnut (csi_csn), and Japanese chestnut (cci_ccn).

### 3.4 Comparative genome structure analysis

The genome structure of Japanese chestnut was well conserved with that of Chinese chestnut (Figure 3), as expected. Interestingly, the *C*. *crenata* genome showed local synteny with the genomes of at least 14 plant species (*Acer yangbiense*, *Citrus sinensis*, *Fragaria vesca*, *Malus* × *domestica*, *Manihot esculenta*, *Populus trichocarpa*, *Prunus avium*, *Prunus dulcis*, *Prunus mume*, *Prunus persica*, *Pyrus betulifolia*, *Theobroma cacao*, *Vitis vinifera*, and *Ziziphus jujuba)* (Figure 3), all of which represent woody tree species belonging to Malpighiales, Malvales, Rosales, Sapindales, and Vitales in rosids^12^.

**Figure 3.**
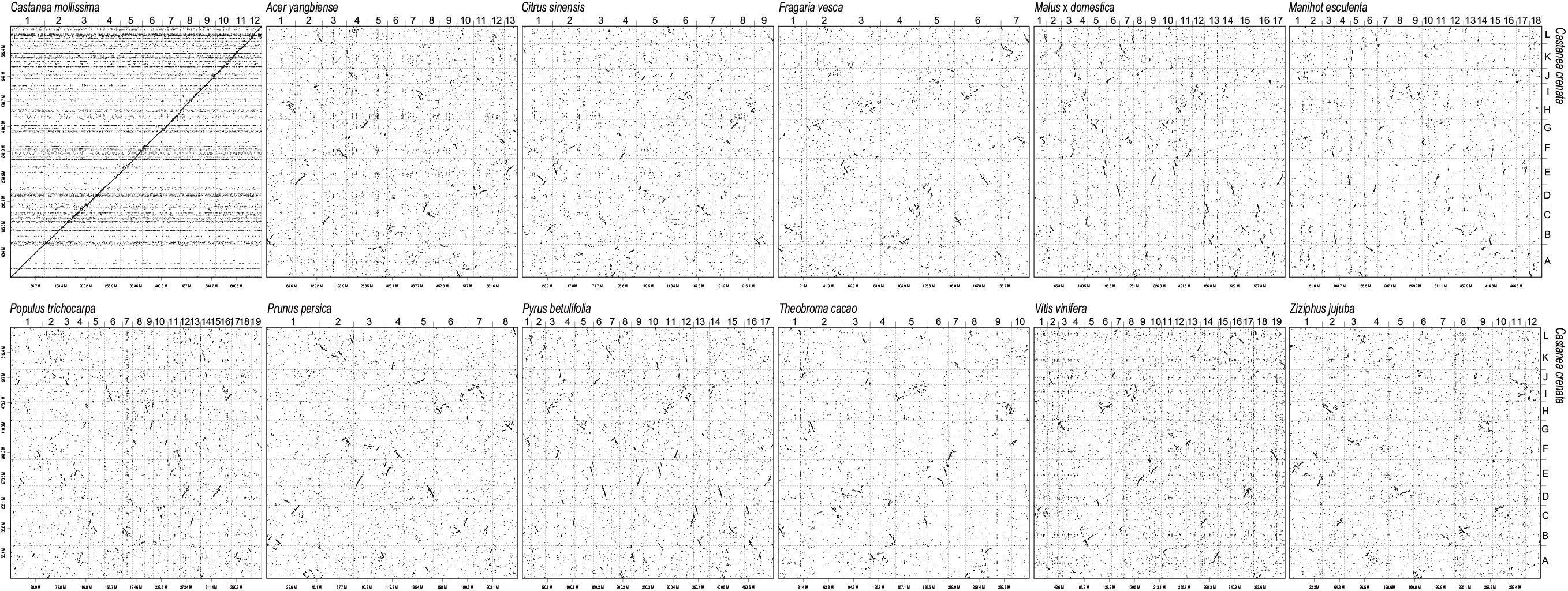
Comparative analysis of the genome sequence and structure of Japanese chestnut and other woody members of rosids. Dots indicate genome sequence and structure similarity among *Castanea mollissima*, *Acer yangbiense*, *Citrus sinensis*, *Fragaria vesca*, *Malus* × *domestica*, *Manihot esculenta*, *Populus trichocarpa*, *Prunus persica*, *Pyrus betulifolia*, *Theobroma cacao*, *Vitis vinifera*, and *Ziziphus jujuba*.

## 4. Conclusion and future perspectives

Here, we report the genome sequence of Japanese chestnut for the first time. Due to the long-sequencing technology and genetic mapping with high-density SNP loci identified using previously published ddRAD-Seq data, 683.8 Mb (94.8%) of the assembled contig sequences (721.2 Mb) were assigned to chromosomes for establishing pseudomolecule sequences for Japanese chestnut (Table 2). Genes predicted based on the pseudomolecule sequences of Japanese chestnut included 96.9% of the core gene set of Embryophyta (Table 2), suggesting that most of all genes were represented in the Japanese chestnut genome assembly. The coverage of pseudomolecule sequences in Japanese chestnut (683.8 Mb) was comparable to that in Chinese chestnut (12 pseudomolecule sequences, 667 Mb). Therefore, the chromosome-level genome assembly of Japanese chestnut generated in this study could be used as a standard reference for genetics, genomics, and breeding of Japanese chestnut and its relatives.

Comparative genome structure analysis showed that the genome structure of Japanese chestnut was quite similar to that of Chinese chestnut (Figure 3). This result suggests that agronomically important genetic loci controlling nut weight and harvesting time^13–15^ could be transferred from Chinese chestnut to Japanese chestnut and vice versa. Chromosomal segments in the Japanese chestnut genome shared synteny with the peach genome^16^ (Figure 3); this relationship has also been reported in Chinese chestnut. More interestingly, regions in the Japanese chestnut genome showed sequence similarity with the genomes of Malpighiales, Malvales, Rosales, Sapindales, and Vitales belonging to woody rosids (Figure 3), but not with Brassicales, Cucurbitales, and Fabales belonging to herbaceous rosids. Genome structure conservation among rosids, e.g., Rosales (sweet cherry, apple, and, Japanese fig) and Malvales (cacao), have been suggested in our previous investigation of 114 plant species^10^. This result suggests that woody members of rosids share a closely related genome structure with each other, although this relationship contradicts that suggested by the APG classification IV^12^. Indeed, none of the plant species in Cucurbitales, which is suggested to be the closest order to Fagales according to APG IV^12^, showed genome structure similarity to Japanese chestnut. Thus, comparative genome structure analysis provides new insights into the evolutionary history of angiosperm.

The genome sequence data generated in this study would enhance the genetics and genomics of Japanese chestnut, in which classical molecular markers, such as simple sequence repeats, previously played a key role in 1) the identification of genetic loci affecting agronomically important traits^4,15,17^; 2) chestnut breeding programs with marker-assisted selection^18^; and 3) the assessment of the genetic structure of cultivated and wild chestnut populations^19^. Furthermore, the highly conserved genome sequence and structure of woody members of rosids might help us better understand their evolutionary history and facilitate the identification of common genetic mechanisms affecting agronomic traits across various orders, families, species, and genera.

## Acknowledgments

We thank Y. Kishida, C. Minami, H. Tsuruoka, and A. Watanabe (Kazusa DNA Research Institute) for providing technical assistance. This work was supported in part by the JSPS KAKENHI (20K15524) and the Kazusa DNA Research Institute Foundation.

## Data availability

Raw sequence reads were deposited into the Sequence Read Archive (SRA) database of the DNA Data Bank of Japan (DDBJ) under accession number DRA012289. Assembled sequences are available at DDBJ under accession numbers BPMU01000001-BPMU01000781. Genome information generated in this study is available at Plant GARDEN (https://plantgarden.jp).

## Conflict of interest

None declared.

## Supplementary data

Supplementary Table S1 Software tools used for genome assembly and gene prediction.

Supplementary Table S2 Genetic map and anchored genome sequences.

Supplementary Table S3 IDs and annotations of predicted genes.

## Notes

### Competing Interest Statement

The authors have declared no competing interest.

